## Supplementary methods and figures for "The Structure and Statistics of Language Jointly Shape Cross-frequency Neural Dynamics During Spoken Language Comprehension"

In this document, we provide additional information and figures that complement the main text of the manuscript. We first elaborate on the computation of the surprisal and entropy features from the GPT2 language model. We then further describe the method used to compute phase-amplitude coupling (PAC) with a more detailed explanation of the estimation of PAC in a feature-dependent manner. Finally, we present additional figures that complement the main text and provide further insights into the results.

### 1 Supplementary Methods

#### Stimulus representation: statistical features

The computation of *surprisal* and *entropy* from the large language model GPT2 [1] needed some care to be adapted to the word-level granularity we chose in our analysis. In fact, GPT2 model computes prediction on *token* which are sub-word units (byte-pair encoding). To compute surprisal and entropy at the word level, we used the following procedure. We first computed the surprisal of each token in the sequence. The conditional next-word probability of a word comprised of several tokens is taken as $P(w_i|w_{i-1}, w_{i-2}, ..., w_1) = \prod_{j=1}^{n} P(t_{ij}|w_{i-1}, ..., w_1)$, where $w_i$ is the word, $t_{ij}$ is the $j^{th}$ token of the $i^{th}$ word, and $n$ is the number of tokens in the word. The surprisal of a word was then computed as $-\sum_j \log(P(t_{ij}|w_{i-1}, ..., w_1))$. That is, the surprisal of a word is taken as the sum of the surprisal of its tokens. On the other hand, the entropy is a measure of the *expected* surprisal at a given word, meaning it summarises the uncertainty of the network in its ability to predict the next word or token. As such, we computed the entropy of word as the entropy of the last token of the word. This indeed entails the network's uncertainty about its prediction once it has "encountered" all token of a given word. The output layer of the network is a *softmax* layer, where each component maps to the probability of the next token. Therefore, the entropy of the word is computed as $-\sum_{j=i}^{N} P(t_i|t_{i-1}..., t_1) \log(P(t_i|t_{i-1}, ..., t_1))$, where we sum over the entire vocabulary of tokens (of length $N$).

#### A note on TRF-based PAC

Several methods have been proposed to estimate the Phase-amplitude coupling (PAC) between two signals. Notably, we can mention the modulation index [2], the Kullback-Leibler divergence [3], the phase-locking value [4] and normalised version of Canolty *et al.* [2]'s modulation index: the direct PAC estimate from Özkurt & Schnitzler [5]. However, these methods are based on trial averaging and thus are not able to disentangle the effect of different continuous experimental covariate on the time-resolved PAC. To address this issue, we propose a method that is directly inspired by the analogy between event-related potential computation (averaging across trials) and the time-resolved regression analysis such as temporal response functions (TRF) [6–8]. The idea is simply to estimate the PAC by modelling the complex phase-amplitude signal –built by multiplying the low-frequency phasor (e.g. for the delta-band: $e^{i\phi_\delta(t)}$) with the power time series at high-frequencies (e.g., for the beta-band: $r_\beta(t)$)– as the convolution of a kernel (which will become our PAC estimate) with the stimulus continuous regressor time-series. This is done per sensor or source location. The kernel is then estimated by regularized regression (ridge regression). The model equation reads, for a single channel:

$$r_\beta(t)e^{i\phi_\delta(t)} = \sum_{j=1}^{N_{feat}} (k_j * x_j)(t) + \epsilon = \sum_{j=1}^{N_{feat}} \sum_{\tau=1}^{\tau_{max}} k_j(\tau)x_j(t-\tau) + \epsilon \tag{1}$$

Where $r_\beta(t)$ is the power time series at the beta frequency band, $e^{\phi_\delta(t)}$ is the phase time series at the delta frequency band, and $x_j(t)$ is the $j^{th}$ feature time series. We thus recover a different time-resolved signal $k_j(\tau)$ for each feature $j$ and each time lag $\tau$. The PAC estimate is then the absolute value of the time-resolved kernel $|k_j(\tau)|$. This method allows us to estimate the PAC in a feature-dependent manner, which is crucial to disentangle the effect of different linguistic features on the time-resolved PAC. We can then compute the PAC for each feature and each time lag, and test the significance of the PAC estimate against a baseline (e.g., the PAC estimate when all linguistic features are nullified).

As such, we assimilate our estimate of PAC to the *modulation index* defined by Canolty *et al.* [2]. This requires to compute the coefficient on surrogate data in order to test for significance of the estimated PAC. In our analysis, we resort to the comparison with null models where the feature vaues for which the PAC is computed has been shuffled. However, we will review the limitation of this method and possible extensions to be investigated in the future. One drawback of this estimate comes from the possible bias by both time-locked evoked activity and power bias. We can see in figure 1 that the delta-beta PAC is not only driven by time-locked phase alignment in the delta-band, as we derive different PAC from the same-low frequency phasor (delta-beta and delta-gamma).

We are not totally exempt of the putative bias from power modulation though. Following Penny *et al.* [4] and Voytek *et al.* [9], a solution could be to compute the phase-locking value between the phase of the low-frequency signal and the phase of the amplitude in the high-frequency band. As such, we could also compute a TRF-based PAC following the phase-locking value defined by Penny *et al.* [4] simply by replacing $r_\beta(t)e^{i\phi_\delta(t)}$ in equation (1) by $e^{i(\phi_\delta(t)-\phi_{r_\beta}(t))}$. Where $\phi_{r_\beta}(t)$ is the phase of the high-frequency amplitude signal. Another possibility is to directly normalise our PAC with TRF-based estimate of feature-based power time course (as in Özkurt & Schnitzler [5]). In other words we could compute the PAC as:

$$\frac{|k_j(t)|}{TRF_{j;\beta}(t)} \tag{2}$$

Where $TRF_{j;\beta}(t)$ is the TRF-based estimate of the *power* time course in the beta frequency band. To conclude, except from the information-theoretic estimation proposed by Tort *et al.* [3], other PAC estimation ([2, 4, 5]) methods can be ported to this TRF-based PAC estimation. Further investigation is required to thoroughly compare the advantages and limitations of each approach. This is left for future work.

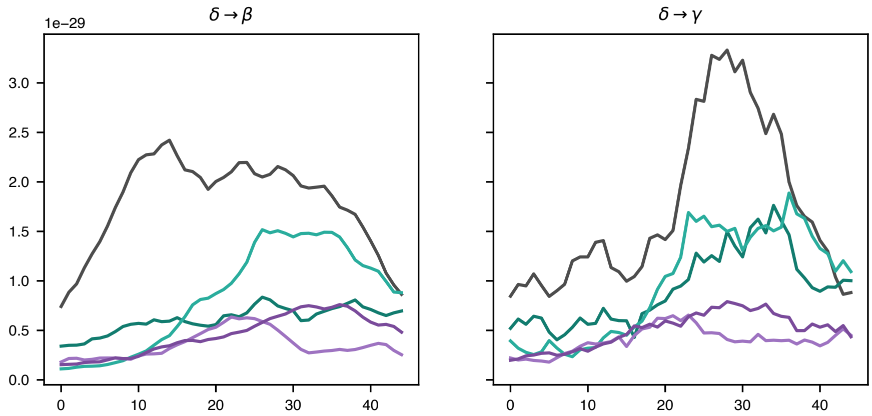


**Fig. 1**: As an example, we show here both delta-beta and delta-gamma PAC for all features except acoustic envelope. This highlights the specificity of each frequency bands, noting that the delta-beta PAC cannot be only driven by time-locked phase alignment in the delta-band.

### 2 Supplementary Figures

We present here additional figures that complement the main text and provide further insights into the results. First, we present the time-resolved score for individual feature in the delta-band. This helps us to better understand the origin of the effect at negative lags observed in Figure 3 of the main manuscript. We then show the time course of the Phase Amplitude Coupling (PAC) coefficients time series as computed for all features, including our acoustic control, as well as an example of the spatio-temporal dynamics of those PAC coefficients in source space.

#### Time-resolved TRF score

The main manuscript shows significant improvement of Rule-based features, and marginal improvement of Statistical features, in the TRF model at negative lags (see Figure 3, panel **c**). We further investigate the time-resolved scoring of TRFs in figure 2. The TRF for each feature are computed separately, using only a short window of lags around the central time point. Then, we compared the score of those models with a model where the extra linguistic feature was shuffled. This provides us with a relative score increment. We observe that the significant improvement in the model at negative lags is driven by the entropy and close features. Importantly, causal filters were used to avoid any anti-causal temporal leakage of information. We note that the improvement of score for entropy and close features at negative lags is not anti-causal. Indeed, the entropy at a given word does *not* depend on the current word heard but only on previous ones. Therefore, the entropy value at a given word can be made accessible to brain process at earlier lags. This is in line with the idea that the brain can anticipate the upcoming words based on the statistics of the language. The close feature, on the other hand, is a proxy for the integration of words into larger syntactic units. We argue (in the manuscript) that the brain indeed anticipates upcoming words which are likely to *close* a larger number of syntactic constituents. More precisely, this shows that it anticipates the structure itself, not necessarily the semantic content of the upcoming words.

#### PAC time course for delta-beta and theta-gamma frequency bands

We show the time course of the PAC coefficients for every feature of our complete model, including the acoustic envelope that was used as a control, in figure 3. We observe a strong theta-gamma coupling in relation to the acoustic envelope (figure 4b). This has been previously observed and suggested as a segmentation mechanism for the acoustic stream of speech [10–12].

Moreover, as an extra control to verify that the resulting PAC where not driven only by the acoustic properties of the signal, we ran our TRF-based PAC analysis for the French condition. We show the time course of the PAC coefficients for the French condition in figure 4. We observe that

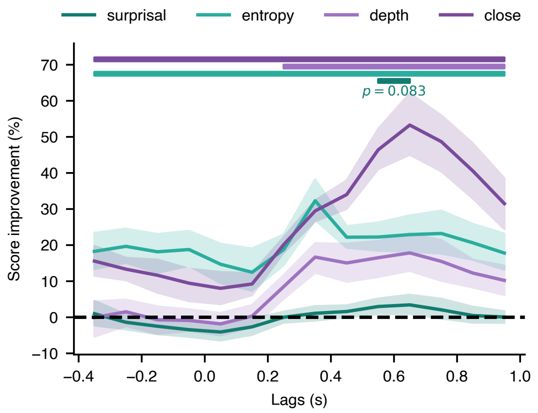


**Fig. 2**: Time resolved scoring of TRFs. Each line corresponds to a different model and each point to a different time segment. Models are computed from a TRF for which we only took a short window (100ms) of lags around the central time point. Models are here built with only word onset and an extra linguistic feature (either Surprisal, Entropy, Depth, or Close). We then compared the score of those model with one where the extra linguistic feature was shuffled to compute the relative score increment (y-axis). This is similar to the analysis of the main manuscript done in figure 3 panel **c**, however without grouping the features per category (statistical or syntactic). We observe that the significant improvement in the model at negative lags is only driven by the entropy and close features. Note that causal filters were used to avoid any anti-causal temporal leakage of information. Top bars indicate significant time for scores against each shuffled model (Wilcoxon signed-rank test, $p<0.05$, except for Surprisal which is marginal, FDR corrected).

the PAC coefficients are not significant for the for linguistic features, only the acoustic features (envelope and word onsets) generate a coupling between low-frequency phase and high-frequency amplitude.

We show as well a more detailed example of how the spatio-temporal coefficients project onto the source space models in figure 5.

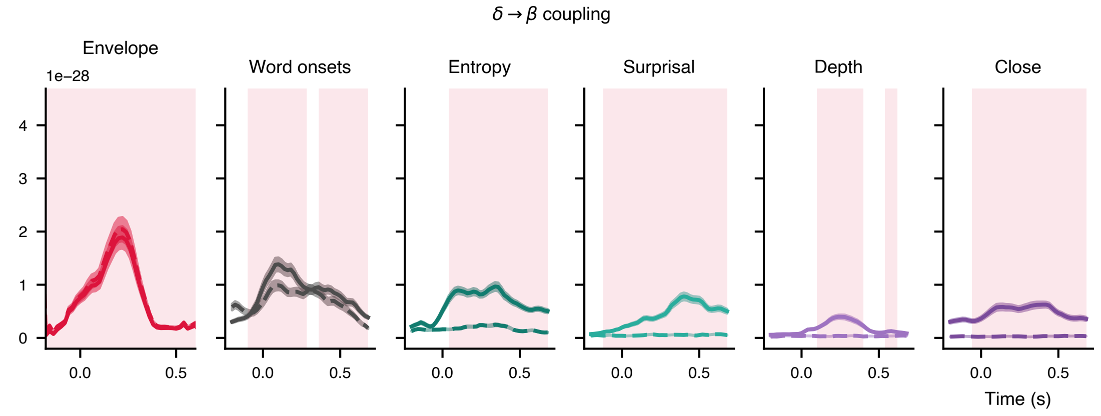


(a) Delta-beta PAC

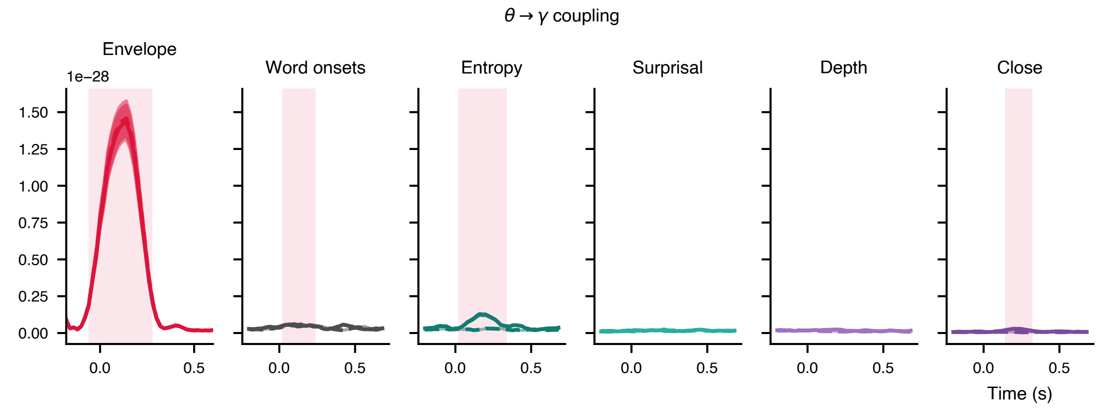


(b) Theta-gamma PAC

**Fig. 3**: Time course of the PAC coefficients for the delta-beta and theta-gamma frequency bands. The top row shows the PAC coefficients for the delta-beta frequency bands, while the bottom row shows the PAC coefficients for the theta-gamma frequency bands. The columns correspond to the different features (Entropy, Surprisal, Depth and Close). We show the global field power across all sensors as a summary statistics of PAC. Shaded areas represent significant cluster for a temporal cluster-based permutation test against baseline (negative lags).

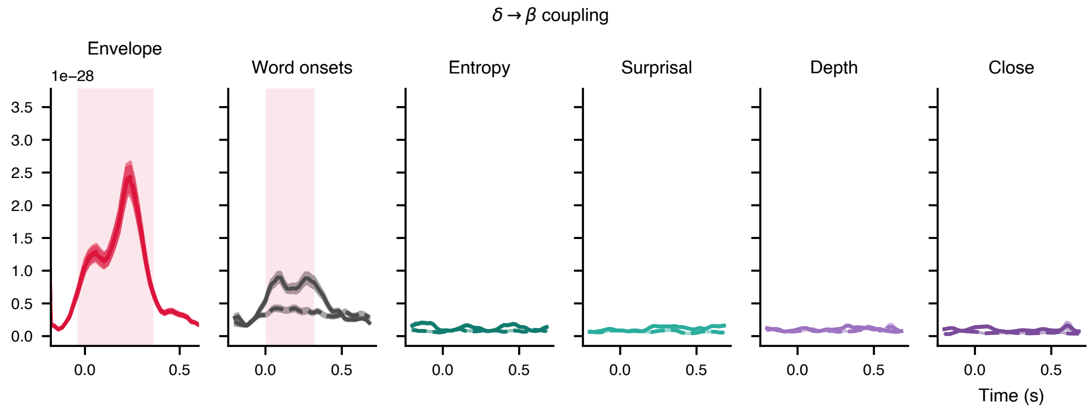


(a) Delta-beta PAC for the French condition

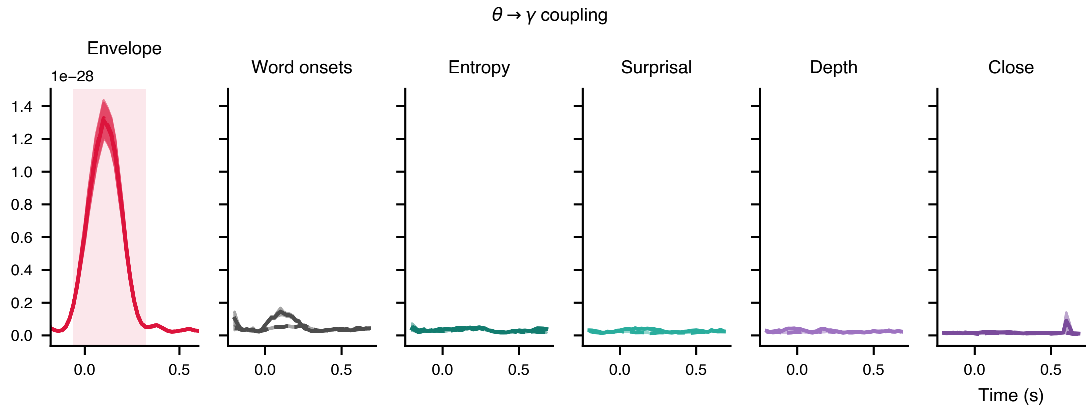


(b) Theta-gamma PAC for the French condition

**Fig. 4**: Time course of the PAC coefficients for the delta-beta and theta-gamma frequency bands computed on the French story parts. Same layout as in figure 3

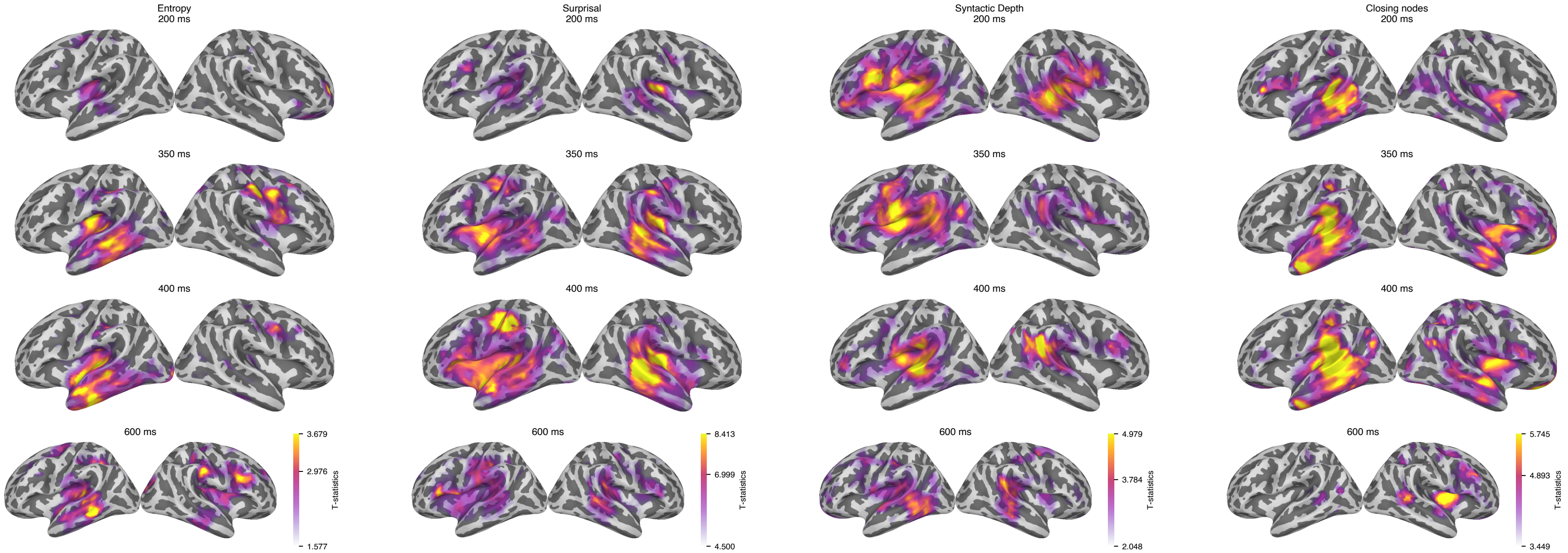


**Fig. 5**: Example of the spatio-temporal development of each feature-PAC time course for the delta-beta couplings. The four row correspond to times: 200ms, 350ms, 400ms and 600ms respectively, while the columns correspond to the different features (Entropy, Surprisal, Depth and Close). The colour scale is the same for all plots and represent the t-statistics against the Base model (in which all linguistic features are *nullified*).

#### References


1. Radford, A., Wu, J., Child, R., Luan, D., Amodei, D. & Sutskever, I. Language models are unsupervised multitask learners. *OpenAI Blog* **1,** 9 (2019).
2. Canolty, R. T. *et al.* High gamma power is phase-locked to theta oscillations in human neocortex. *science* **313,** 1626–1628 (2006).
3. Tort, A. B. L., Komorowski, R., Eichenbaum, H. & Kopell, N. Measuring Phase-Amplitude Coupling Between Neuronal Oscillations of Different Frequencies. *Journal of Neurophysiology* **104.** PMID: 20463205, 1195–1210. eprint: https://doi.org/10.1152/jn.00106.2010. https://doi.org/10.1152/jn.00106.2010 (2010).
4. Penny, W., Duzel, E., Miller, K. & Ojemann, J. Testing for nested oscillation. *Journal of Neuroscience Methods* **174,** 50–61. ISSN: 0165-0270. https://www.sciencedirect.com/science/article/pii/S0165027008003816 (2008).
5. Özkurt, T. E. & Schnitzler, A. A critical note on the definition of phase–amplitude cross-frequency coupling. *Journal of Neuroscience Methods* **201,** 438–443. ISSN: 0165-0270. https://www.sciencedirect.com/science/article/pii/S0165027011004730 (2011).
6. Di Liberto, G. M. *et al.* Low-Frequency Cortical Entrainment to Speech Reflects Phoneme-Level Processing. *Current Biology* **25,** 2457–2465. ISSN: 09609822. http://www.sciencedirect.com/science/article/pii/S0960982215010015%20http://www.ncbi.nlm.nih.gov/pubmed/26412129%20https://linkinghub.elsevier.com/retrieve/pii/S0960982215010015 (Sept. 2015).
7. Broderick, M. P., Anderson, A. J. & Lalor, E. C. Semantic Context Enhances the Early Auditory Encoding of Natural Speech. *The Journal of neuroscience : the official journal of the Society for Neuroscience* **39.** ISSN: 15292401 (2019).
8. Weissbart, H., Kandylaki, K. D. & Reichenbach, T. Cortical Tracking of Surprisal during Continuous Speech Comprehension. *Journal of Cognitive Neuroscience* **32,** 155–166. ISSN: 0898-929X. https://www.mitpressjournals.org/doi/abs/10.1162/jocn%7B%5C_%7Da%7B%5C_%7D01467 (Jan. 2020).
9. Voytek, B., Canolty, R., Shestyuk, A., Crone, N., Parvizi, J. & Knight, R. Shifts in Gamma Phase–Amplitude Coupling Frequency from Theta to Alpha Over Posterior Cortex During Visual Tasks. *Frontiers in Human Neuroscience* **4.** ISSN: 1662-5161. https://www.frontiersin.org/articles/10.3389/fnhum.2010.00191 (2010).
10. Giraud, A.-L. & Poeppel, D. Cortical oscillations and speech processing: emerging computational principles and operations. *Nature Neuroscience* **15,** 511–517. ISSN: 1546-1726. http://www.ncbi.nlm.nih.gov/pubmed/22426255%20http://www.nature.com/articles/nn.3063 (Apr. 2012).
11. Fontolan, L., Morillon, B., Liegeois-Chauvel, C. & Giraud, A.-L. The contribution of frequency-specific activity to hierarchical information processing in the human auditory cortex. *Nature Communications* **5,** 4694. ISSN: 2041-1723. https://doi.org/10.1038/ncomms5694 (Sept. 2014).
12. Hovsepyan, S., Olasagasti, I. & Giraud, A.-L. Rhythmic modulation of prediction errors: A top-down gating role for the beta-range in speech processing. *PLOS Computational Biology* **19,** 1–29. https://doi.org/10.1371/journal.pcbi.1011595 (Nov. 2023).
